# Acute effect of transcranial direct current stimulation (tDCS) on postural control of trained athletes: a randomized controlled trial

**DOI:** 10.1101/2023.05.17.541119

**Authors:** Mary Giancatarina, Yohan Grandperrin, Magali Nicolier, Philippe Gimenez, Chrystelle Vidal, Gregory Tio, Emmanuel Haffen, Djamila Bennabi, Sidney Grosprêtre

## Abstract

Transcranial direct current stimulation (tDCS) is used to modulate the brain function in targeted brain areas, and can acutely modulate motor control, such as postural control. While the acute effect of tDCS is well documented on patients, little is still known whether tDCS can alter the motor control of healthy populations with an already high level of motor skills. This study aimed to assess the acute effect of tDCS on postural control of trained athletes. Eighteen parkour practitioners, known for their good balance abilities, were tested on three occasions in the laboratory for each stimulation condition (2 mA ; 20 minutes) – primary motor cortex (M1), prefrontal cortex (dlPFC) and sham (placebo). Postural control was evaluated PRE and POST each stimulation by measuring Center of Pressure (CoP) displacements on a force platform during static conditions (bipedal and unipedal stance). Following M1 stimulation, significant decreases were observed in CoP area in unipedal (P=0.003) and bipedal (P<0.001) stances. As well, the CoP total length was significantly reduced in bipodal (P=0.005) as well as in unipedal stance (P<0.001), only after M1 stimulation. Relative pre-post changes observed after M1 stimulation were negatively correlated to experience in parkour only after unipedal stance (r=0.715, P<0.001), meaning that the more participants were trained the less tDCS was effective. No significant changes were noticed after sham and dlPFC stimulation. These results suggested that the modulation of gait performance in athletes following an acute intervention of tDCS is specific to the targeted brain region, and that more complex postures (such as unipedal stance) were more sensitive to the effect of tDCS according to the level of practice.

## INTRODUCTION

Postural control requires the interaction of several brain regions, including the motor and pre-motor cortex, the sensorimotor cortex, the basal ganglia, the thalamus, the fronto-parietal regions and the cerebellum [1]. Among the methods that improve balance, non-invasive brain stimulation has been suggested as a good alternative to improve motor skills or optimize rehabilitation programs [2]. Transcranial direct current stimulation (tDCS) consists of delivering a constant low intensity current (1 to 4 mA) evoked by an electrode (the anode) and directed towards a second electrode (the cathode). Through this process, tDCS is able to modulate neural activity of the cortical crossed areas [3]. The literature about tDCS and postural control has mainly focused on the primary motor cortex, showing improvement either during [4] or after the stimulation [5,6]. It is suggested that modulating M1 excitability might improve balance by modulating cortico-muscular coherence in ankle muscles [4,7]. However, another brain region is often targeted in tDCS literature: the left dorsolateral prefrontal cortex (dlPFC). So far, this brain region is mostly targeted for the improvement of cognitive functions such as attention [8] or memory [9]. The dlPFC is also a frequent target to evaluate whether tDCS can improve motor performance, mostly related to whole-body and/or single joint endurance [10,11], but also to fine motor skills during reaching or pointing tasks [12,13]. There already exist some clues in the literature, although much less than for M1, that stimulation of the dlPFC also appears likely to improve postural control in young recreationally active adults [14]. Indeed, several recent structural and functional neuroimaging studies showed that dlPFC is involved in the control of standing and locomotion [15].

Most previous works on the tDCS effect on postural control included untrained subjects with no specific skills in balance, questioning if tDCS could work on pre-existant circuitry or if a ceiling effect could be observed on more experienced participants. It is well known that sport practice leads to refine postural control [16]. In the present study, we aimed to include athletes that are particularly trained regarding balance – parkour practitioners [17]. Parkour is a modern physical activity which consists in overcoming various obstacles from the urban landscape, such as fences, benches, or walls [18].

Proprioception and balance are therefore widely used by Parkour practitioners (also called traceurs), leading them to have a better perception of their body in space in comparison to recreationally active individuals [17]. This last point also allows to counterbalance the possible lack of visual information and justifies that traceurs have a significantly lower Centre of Pressure (CoP) surface area with their eyes closed than recreationally active subjects [17]. Besides, Parkour offers a particular dynamic postural stability – especially when performing precision landings [19] for which practitioners develop a more stable compensatory movement to counteract the impact at landing, with a lower excursion of the CoP than sedentary subjects. We previously demonstrated that tDCS acutely applied over M1 led to a significant improvement of jump abilities of traceurs [12]. Conversely, tDCS applied over dlPFC did not lead to jump improvement but refined the management of fine motor tasks, i.e. accurate pointing tasks [12]. Therefore, managing posture is at the interface of a whole-body performance such as jumping and a fine motor control strategy showing that either dlPFC or M1 stimulation could possibly modify balance ability in parkour practitioners.

In this context, we aimed to assess the acute effect of tDCS applied on two brain regions classically stimulated in the literature (M1 and the left dlPFC) on postural control of athletes with well-known balance abilities. Mostly because of a potential ceiling effect, the effect of tDCS might be different in trained participants than in recreationally active participants as studied in previous literature, and particularly the brain region of interest. We therefore conducted a double-blinded, sham-controlled trial performed by recruiting participants with various experience in Parkour, from beginners to well-trained athletes. It was suggested that tDCS effects can be widely influenced by many factors, including the type and amount of previous sport practice [20]. Given that this population present pre-cabled cortical circuits to regulate balance, we hypothesized that tDCS may have a significant effect on postural control of Parkour athletes. However, a ceiling effect could be expected on the most experienced individuals as suggested in previous experiments on power-lifters [21] or endurance athletes [22]. Then, we also hypothesized that tDCS might have lower effect in the most trained individuals.

## MATERIAL AND METHODS

### Participants

Eighteen healthy young males (age: 22.6 ± 5.7 years old; height: 180 ± 5.7 cm; weight: 74.5 ± 7.8 kg) gave their written informed consent to participate in the present study. Participants reported a regular practice of Parkour, at a wide range of expertise. Their total training volume (in hours), estimated with their experience (in years) and training frequency (in hours/week) has been evaluated and ranged from 156 to 10,608 hours (mean: 4343 ± 3508). Research protocol was approved by the regional ethic committee (CPP-Est-IV) under the number 18/47, registered on ClinicalTrials.org (NCT03937115), and conducted in accordance with the last version of the Declaration of Helsinki.

### Experimental set-up and procedures

This study was a double-blinded, randomized, sham-controlled trial to determine the effects of tDCS applied over M1 or dlPFC on static and dynamic balance of Parkour practitioners. The protocol was carried out in 3 experimental sessions separated by at least 48h. The following conditions were performed randomly – anodal tDCS over right M1, anodal tDCS over dlPFC and sham stimulation. To ensure the blinding of the type of tDCS applied toward participants, participants were told that any of the three experimental conditions could be a placebo session, independently of the electrode placement. Participants were asked after each session to report if they felt the stimulation was active or placebo. Most subjects (55.6%) thought they had received active stimulation during the placebo session. No difference has been reported in adverse events experienced (tingling, itching, etc.) between the placebo and the active sessions. During each session, subjects performed postural tasks on a force plate before and after tDCS intervention. They executed a bipedal and unipedal static posture before and after tDCS conditions. PRE and POST tests were randomized.

### Assessment of static posture

Static posture was appraised by means of two experimental conditions – bipedal and unipedal stance, performed on a force plate (Kistler Instrument Corp., Winterthur, Switzerland). The force plate allowed continuous recording of CoP displacements (sampling frequency: 1000 Hz) in both medio-lateral and postero-anterior axis. Bipedal stance was assessed by asking participants to stand still on the force plate, feet shoulder-width apart and arms relaxed. Unipedal stance was evaluated by asking participants to stand still on the left foot by bending the knee of the right leg. Contact with the supporting leg was not allowed. Foot placement and spacing was measured during the first session to be reproduced in the two others. In the two postural conditions, 30 seconds recording were performed, during which participants were asked to keep quiet, eyes open. The last 20 seconds were taken for analysis.

### Transcranial Direct Current Stimulation

tDCS was delivered by a neurostimulation device (StarStim®, Neuroelectrics©, Barcelona, Spain) adapted to double-blind procedure. It was transmitted by two saline-soaked synthetic sponge electrodes (Sponstim®, 25cm^2^) placed on the scalp or shoulder. Every participant agreed to undergo three 20-minute sessions of tDCS (two actives and one sham), separated by 48 hours. Electrodes placement was carried out using the international EEG 10/20 system. The experiment included three stimulation sequences: (i) anode facing the dlPFC (F3 position) and cathode facing the right supra-orbital region (AF8 position), (ii) anode over the dlPFC (F3) and cathode facing the right supra-orbital region (AF8), which is related to sham condition, and (iii) anode facing the right M1 (FC2 position) and cathode placed on the left shoulder. Sequence order was randomized by computer beforehand. During active stimulation conditions, a 2 mA current was generated. The sham condition corresponds to a gradual increase in the intensity of the current during the first 30 seconds up to 2 mA (ramp-up), as recommended by previous literature (Gandiga et al., 2006; Moreira et al., 2021; Swann et al., 2015)

### Data analysis

In static conditions, i.e. bipedal and unipedal upright standing, CoP characteristics were analyzed taking the 20 seconds trial as a whole. The force-plate represents a two-dimensional plane in which the x-axis represents the postero-anterior axis and the y axis the medio-lateral axis, since participants were always oriented in the same direction. The total sway path (length of CoP displacement) has been determined as the total distance covered by the CoP during the 20 seconds of interest. The area of CoP displacements was also analyzed as the area of the ellipse that includes 90% of CoP points for 20 seconds. Postero-anterior and medio-lateral amplitudes were determined as the maximal amplitude (maximal value – minimal value) covered by the CoP in the ellipse area in the x-axis and y-axis, respectively.

The relationship between these CoP variables and training volume of the participants (in hours, see “participants” section) were evaluated by determining the coefficient of correlation (Pearson’s correlation) to test the effect of experience on postural control.

The relative changes in CoP variables with the different tDCS intervention were determined in percentage by the following formula: [(POST-PRE)/PRE]*100. These relative changes plotted against training volume or initial performance (averaged PRE value).

### Statistical analysis

All data are presented as the mean ± standard deviation (S.D.). Each test (unipedal, bipedal) was considered separately. The normality of the data sets was verified by the Shapiro-Wilk test, and variance homogeneity by the Levene test. Two-way repeated measures ANOVA were used to assess differences between the 3 conditions with factors “time” (pre, post) and “tDCS intervention” (M1, dlPFC, SHAM). A one-way repeated measures ANOVA was performed on the relative changes, with factor “tDCS intervention” (M1, dlPFC, SHAM). Post-hoc tests were performed with Bonferroni test. Relationships between the several variables were assessed through Pearson’s coefficient of correlation. To account for session order effect (since tDCS conditions were performed randomly), two-way repeated measures ANOVA were also used to assess differences between the 3 conditions in their chronological order with factors “time” (pre, post) and “number of the session” (first, second, third). Effects size are indicated as the partial eta square (η^2^p), Statistical analysis was performed using STATISTICA (10.0 version, Statsoft, Tulsa, Okhlaoma, USA). The level of statistical significance was set at P < 0.05.

## RESULTS

Regarding a potential confounding factor due to the three separate days of testing, we carefully checked the baseline measurement during each session. No difference was found in any parameter regarding differences between the PRE data of each experimental session. Similarly, no session-order effect was found when analyzing data chronologically, regardless of the tDCS intervention.

No significant changes in CoP length and area have been observed for both sham and dlPFC conditions, in both bidepal and unipedal stance (figure 1A, B, C D). Following M1 stimulation, significant decreases were observed in CoP length from PRE to POST in bipedal stance (P=0.004) and unipedal stance (P=0.003). As well, significant decreases were also observed in CoP area (P<0.001 and P=0.002 for bidepal and unipedal stances, respectively). All statistic results are presented in table 1.

**Figure 1.**
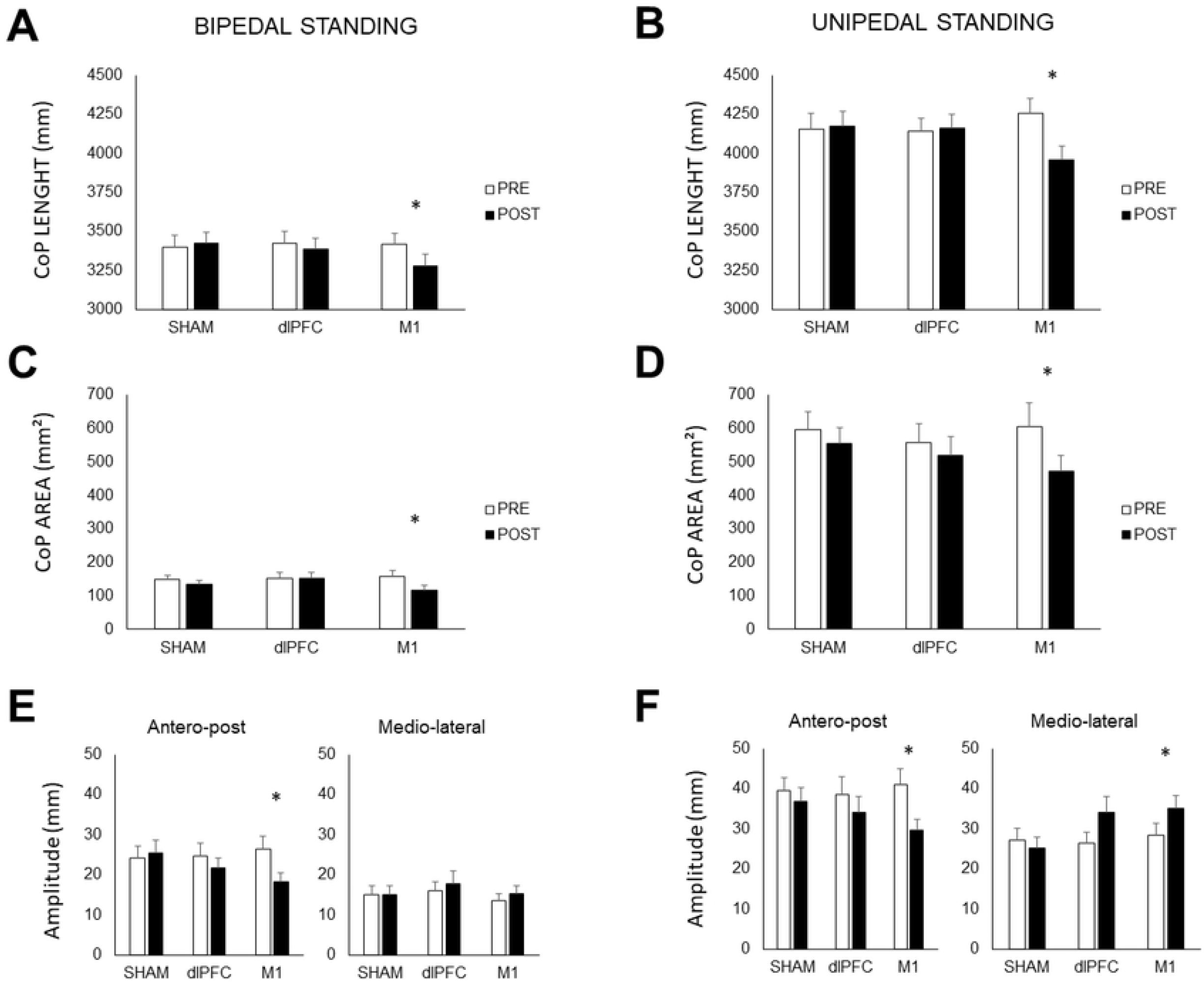
Evolution of Center of Pressure (CoP) characteristics and area before (PRE) and after (POST) each tDCS session. Data are displayed as mean ± SEM. SHAM: placebo stimulation. dlPFC: dorsolateral prefrontal cortex. M1: primary motor cortex. CoP total length in bipedal standing (A.) and unipedal standing (B.). CoP ellipse area in bipedal posture (C) and unipedal posture (D). Postero-anterior (x axis) and medio-lateral (y-axis) amplitude of CoP displacement were also quantified, in bipedal stance (E) and unipedal stance (F). *: significant pre-post difference

**Table 1.** Data and statistical results of bipedal and unipedal stance following tDCS conditions.

| CoP characteristics |  | SHAM |  | dIPFC |  | M1 |  | ANOVA interaction<br>(intervention x time) |  |  |
| --- | --- | --- | --- | --- | --- | --- | --- | --- | --- | --- |
|  |  | PRE | POST | PRE | POST | PRE | POST | F (2,34) | P | η²p |
| BIPEDAL |  |  |  |  |  |  |  |  |  |  |
| Total Lenght (mm²) | MEAN | 3398,88 | 3424,31 | 3425,74 | 3387,09 | 3416,58 | 3280,59 | 5,738 | 0,007 | 0,252 |
|  | SD | 317,19 | 306,48 | 315,00 | 289,72 | 295,37 | 306,23 |  |  |  |
| Postero-anterior amplitude (mm) | MEAN | 24,27 | 25,45 | 24,79 | 21,63 | 26,32 | 18,34 | 7,523 | 0,002 | 0,307 |
|  | SD | 11,82 | 13,40 | 12,89 | 11,33 | 14,51 | 9,35 |  |  |  |
| Medio-lateral amplitude (mm) | MEAN | 15,04 | 14,97 | 15,93 | 17,88 | 13,46 | 15,24 | 0,372 | 0,692 | 0,021 |
|  | SD | 9,44 | 10,23 | 9,50 | 12,79 | 7,52 | 8,83 |  |  |  |
| CoP ellipse (cm²) | MEAN | 149,04 | 135,65 | 151,25 | 150,85 | 157,45 | 117,59 | 8,989 | 0,001 | 0,346 |
|  | SD | 51,16 | 48,68 | 78,20 | 78,08 | 74,04 | 59,82 |  |  |  |
| UNIPEDAL |  |  |  |  |  |  |  |  |  |  |
| Total Lenght (mm²) | MEAN | 4155,78 | 4174,93 | 4141,35 | 4159,01 | 4259,57 | 3846,46 | 11,765 | 0,000 | 0,409 |
|  | SD | 416,28 | 410,63 | 342,91 | 385,84 | 398,36 | 468,96 |  |  |  |
| Postero-anterior amplitude (mm) | MEAN | 39,55 | 36,70 | 38,53 | 34,14 | 41,06 | 29,73 | 3,403 | 0,045 | 0,167 |
|  | SD | 13,74 | 15,50 | 18,67 | 16,52 | 16,49 | 10,80 |  |  |  |
| Medio-lateral amplitude (mm) | MEAN | 27,13 | 25,13 | 26,47 | 34,03 | 28,37 | 35,00 | 5,622 | 0,008 | 0,249 |
|  | SD | 12,31 | 11,86 | 11,28 | 17,26 | 12,55 | 14,44 |  |  |  |
| CoP ellipse (cm²) | MEAN | 596,74 | 555,47 | 558,79 | 520,67 | 607,09 | 451,11 | 3,202 | 0,053 | 0,159 |
|  | SD | 220,84 | 204,01 | 231,32 | 228,61 | 297,87 | 173,91 |  |  |  |

In bidepal stance, a significant decrease from PRE to POST has been observed after M1 stimulation in postero-anterior amplitude of CoP displacement (P=0.043), while no change has been observed in mediolateral axis (figure 1E). In unipedal stance, significant decrease of postero-anterior amplitude (P=0.003) has been observed, while mediolateral amplitude decreased (P=0.005), both after M1 stimulation only (figure 1F). No significant changes in amplitudes have been observed from PRE to POST in both sham and dlPFC conditions.

It can be noticed that relationships have been established between CoP characteristics and level of expertise of the participants. First, both CoP lengths and area were significantly and negatively correlated to the training volume, being lowered as the training volume was higher (figure 2A, B, C, D). Interestingly, the relative PRE-POST changes in CoP lenght induced by M1 stimulation was significantly correlated to the initial CoP length (PRE-data) for unipedal stance only (figure 2F).

**Figure 2.**
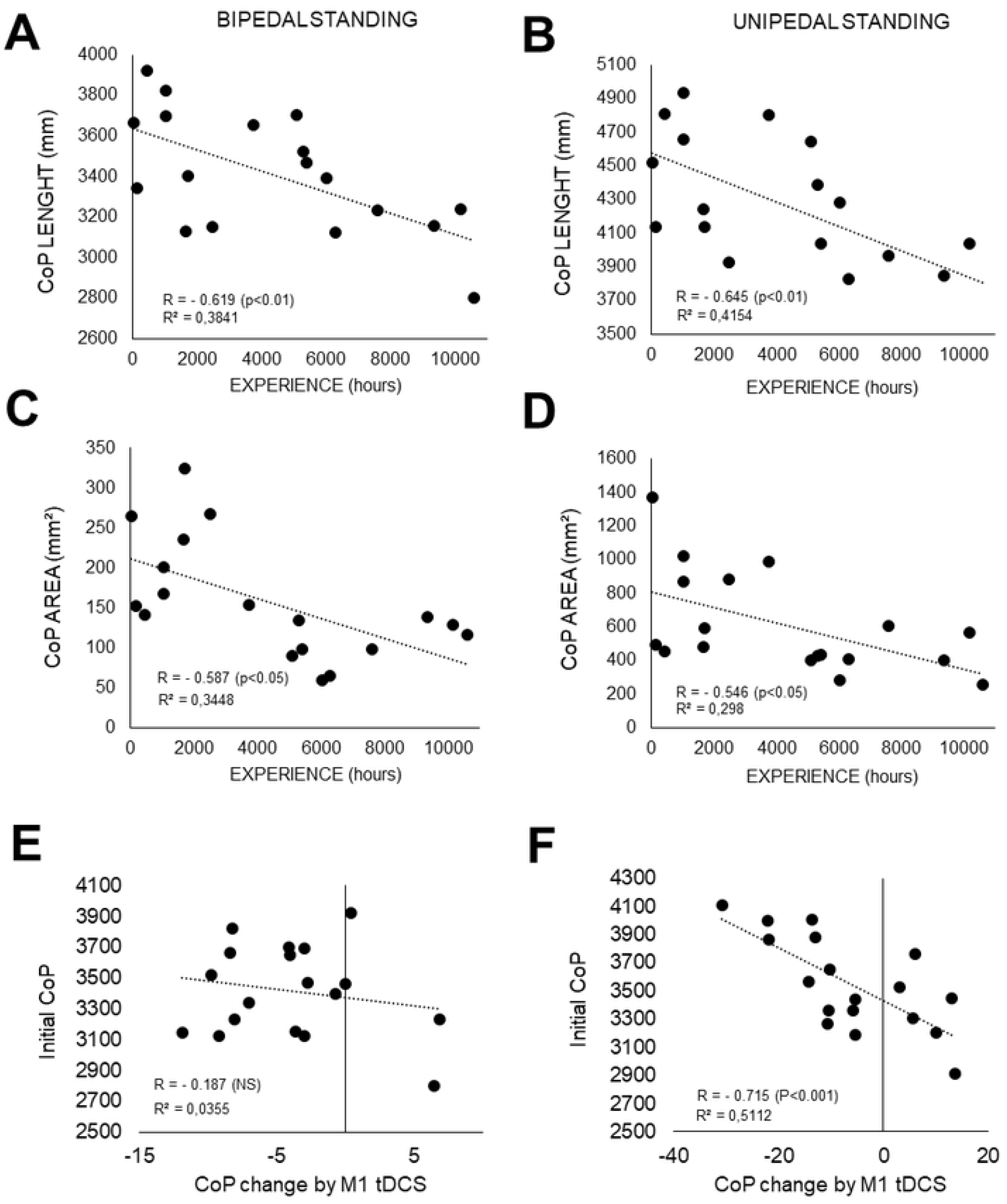
Relationships between experience and CoP characteristics during static stance. A, C, E: bidepal recordings. Initial CoP performances represent the averaged pre value by participants, before any tDCS intervention. B, D, F: unipedal recordings. In E and F, the relative pre-post evolution of CoP with M1 tDCS are plotted against the initial CoP lenght value for bipedal and unipedal recordings, respectively. The relative changes in CoP variables with the M1 tDCS intervention is determined in percentage by the following formula: [(pre-post)/pre]*100).

## DISCUSSION

The effect of two different montage of tDCS on postural control of Parkour athletes has been monitored by analyzing the CoP characteristics in static and dynamic tasks. An acute session of tDCS applied to M1 led to significant reduction of CoP movements for unipedal and bipedal stances while dlPFC or SHAM stimulation did not. This decrease was negatively correlated to the experience of the participants, i.e. lower for the most trained athletes.

### Neurophysiological mechanisms

It has been shown that anodal stimulation of dlPFC could lead to an improvement of postural control, in younger healthy subject [14]. The result of the present study did not meet the initial hypothesis since no effect has been found in parkour practitioners. A possible explanation of this lack of results could probably be linked to the convergence of two factors: 1) parkour athletes present already high balance abilities 2) the dlPFC is not the most involved area in posture. Indeed, traceurs demonstrated strong abilities to maintain posture at least in the tested tasks [17], making the room for improvement far less important than in recreationally active participants. Therefore, although the dlPFC do have a role in managing standing balance [15], its role remains minor as compared to other brain region. The dlPFC is known to play a role in motor planning [23], and its stimulation was shown to be effective in modulating accurate pointing performances in parkour practitioners [12]. From the present result, regarding posture it seems that parkour athletes would need a stronger stimulus.

On the contrary and in line with previous results on healthy young recreationally active participants [4,6,24,25], anodal M1 stimulation had a positive effect on balance control. M1 stimulation would be associated with neurophysiological changes over several hours [26] corresponding to an increase in the motor region excitability which would prevent decline in downward neural control. tDCS would also be able to modulate the motor cortical transmitter system by decreasing the activity of the gamma-aminobutyric acid (GABA)-ergic system [27], whose primary function is to inhibit nerve impulses. tDCS applied to M1 would therefore strengthen postural control by its ability to request more neural networks of the sensorimotor system [7,28]. In addition, this trial adopted an extracephalic montage, which maximizes the effect of tDCS over the anode and can imply the modulation of a larger neural path [12], including subcortical structures involved in postural control[28]. Thus, the neural activation of muscles engaged in maintaining balance would be more efficient, which could justify the reduction of CoP displacements for static stance.

However, M1 stimulation did not produce identical results between bipedal and unipedal balance. Even if the decrease of both CoP length and area was significant for both bipedal and unipedal conditions, this was not the case for medio-lateral (ML) displacements, for which the PRE-POST difference was significantly higher in unipedal condition only. More precisely, unipedal stance showed an increase in the ML movements after stimulation, while those made in the postero-anterior (PA) axis were reduced. This result was not expected but could finally attest for a change in postural control strategy [29]. Unipedal balance requiring offsetting movements, decreasing sway on the PA axis could lead to a strong increase of ML oscillations. Unipedal balance involves tightening the calf muscles – sural triceps including gastrocnemii and soleus muscles – to reduce PA displacements or contracting the foot soles and ankles stabilizing muscles, such as the peroneus longus, to limit movements along the ML axis. These two coping strategies do not appear during the bipedal balance.

Considered the reference posture in humans, standing position offers better stability given that the support polygon in which we are involved is larger. Moreover, our results clearly showed that the amplitude of displacements on both axes was less important during bipedal condition (figure 1E, F).These findings agree with some previous research, where larger PA movements were detected when keeping posture became less easy [30]. Bipedal posture may then not be sufficiently sensitive to small variations such as those induced by tDCS, particularly in well-trained athletes. Unipedal standing seemed to represent a more complex task, allowing to discriminate the effect of tDCS. Indeed, for instance the ellipse of CoP displacement was nearly 5 times wider in unipedal stance than in bipedal stance. Stimulating M1 had then a wide field of action to enhance the characteristics of CoP in unilateral task (figure 2F). Motor abilities and particularly postural strategies can become more effective with training and practice [31]. Therefore, even with a good initial performance in balance, tDCS was efficient to modulate unipedal stance in well trained athletes. This possibly means that tDCS similarly affected uni- and bi-pedal strategies.

### tDCS and targeted population

tDCS’s impact varied depending on the level of expertise of the participants. The systematic review of Brachman et al., 2017 concluded that balance training was an effective tool in improving postural control [32]. Holding balance is therefore at the center of training which justifies that the most experienced traceurs showed a lower CoP length and area. Parkour practitioners already demonstrated a significant better postural control performance than sedentary people during landing [19]. The effects of physical activity on postural control depends on the time of practice [33]. Traceurs, therefore, represent a population on which brain stimulation is effective on postural balance through the stimulation of pre-existing neural networks. However, a ceiling effect can be reached for more experienced practitioners. When carefully looking at individual data, it can be observed that for some individuals the CoP length did not decrease after M1 tDCS. Therefore, a limit can be established between those who showed a significant downward modulation of CoP length after tDCS (n=15/18 for bipedal stance and n=12/18 for unipedal stance) for and those who did not (n=3/18 for bipedal stance and n=6/18 for unipedal stance). According to the training volume of these participants, the limit could be established between 6000 hours (for unipedal stance) and 8000 hours of training (for bipedal stance). Therefore, it appears that this ceiling effect appears from these amounts of training, for which the athlete would probably not benefit from this specific tDCS intervention, at least for the tested postural tasks. In fact, the more the subject is trained, the less room there is for improvement. Pushing back the limit of athletes requires more and more effort for trainers, while gains become smaller and smaller as the athletes’ progress. Finally, it can also be noticed that a great inter and intra-subject variability was reported regarding corticospinal excitability modulation following tDCS, which can be linked with a lack of knowledge concerning the current transfer location [34]. Besides participants’ expertise in the applied tasks, the effects of tDCS may also be affected by other intrinsic factors such as individual anatomic variations [35].

### Conclusion

To summarize, the results of our study suggest that offline M1 stimulation with an extracephalic montage was suitable to improve postural control in Parkour practitioners, contrary to dlPFC stimulation. Among other intrinsic factors, we found that the effects of tDCS were correlated with participants’ expertise in Parkour, as expressed by their total training volume. Then, although tDCS seems to be effective in modulating postural control in athletes, this technique may have limitations in the more highly trained individuals.

## ACKNOWLEDGEMENTS

Stimulation material was courtesy of NEURAXESS (NeuroImagerie Fonctionnelle et NeuroStimulation) research center, Besançon. The authors would like to thank the subjects for their time and enthusiasm. The authors are particularly grateful to the French Parkour Federation (Fédération de Parkour, FPK) for its help and support.

